# CRISPRCasIdentifier: Machine learning for accurate identification and classification of CRISPR-Cas systems

**DOI:** 10.1101/817619

**Authors:** Victor A. Padilha, Omer S. Alkhnbashi, Shiraz A. Shah, André C. P. L. F. de Carvalho, Rolf Backofen

## Abstract

CRISPR-Cas genes are extraordinarily diverse and evolve rapidly when compared to other prokaryotic genes. With the rapid increase in newly sequenced archaeal and bacterial genomes, manual identification of CRISPR-Cas systems is no longer viable. Thus, an automated approach is required for advancing our understanding of the evolution and diversity of these systems, and for finding new candidates for genome engineering in eukaryotic models. In this paper, we introduce a holistic strategy that combines regression and classification models for improving the quality of protein cascades, predicting their subtypes, detecting signature genes and extracting potential rules that reveal functional modules for CRISPR.

## INTRODUCTION

CRISPR-Cas systems provide archaea and bacteria with a nucleic acid based adaptive immune system against invading viruses and plasmids. Mechanistically, the immune response can be divided into three stages, namely adaptation, processing and interference, each carried out by different sets of protein complexes (1). The universally conserved proteins Cas1, Cas2 and, optionally Cas4, are responsible for the adaptation stage, when a fragment of invader DNA is excised and stored in the host chromosome as a spacer in the non-coding CRISPR region. The processing and interference stages are much more mechanistically diverse, using different sets of proteins, depending on the type of CRISPR-Cas system. CRISPR-Cas systems are found in many bacteria and most archaea, and have diversified as much as their host organisms (2).

While the mechanistic principles are similar, with spacers comprising templates for synthesis of CRISPR interference RNAs (crRNAs) against the invader, the different types and classes of CRISPR-Cas systems show some important differences. Class 2 systems use a single multi-domain protein for locating and cleaving the re-invading nucleic acid, whereas Class 1 systems employ a large multi-subunit complex for the same purpose. Class 2 systems can be further subdivided into type II, V and VI, which appear to have evolved independently from each other. Thus, the respective Cas9,12 and 13 enzymes that carry out invader cleavage rely on diverse mechanisms involving differing nuclease domains (2, 3, 4).

Class 1 systems, on the other hand, with types I, III and IV, use structurally related proteins to carry out similar functions, although the protein subunits have diverged considerably. Common to all Class I systems is that Cas7 forms a helical backbone that spans the length of the tightly bound crRNA. This backbone is terminated in one end by Cas5, which itself is bound to Cas8 or Cas10 for type I and IV, or type III systems, respectively. Type I systems use Cas8 for the recognition of the protospacer adjacent motif (PAM) (5), which, along with invader crRNA hybridisation, comprises a signal for recruitment of the Cas3 helicase-nuclease protein that subsequently digests the invader chromosome (6). Type III systems, however, use the Cas7 backbone for cleaving invader mRNA while the Cas10 HD nuclease cleaves transcribed DNA (7, 8). Cas10 also synthesises a signaling molecule that recruits additional accessory Cas proteins for other functions, such as cell suicide or activation of other defense systems (8, 9).

The different types of CRISPR-Cas systems are themselves so diverse that each type can be further subdivided into several subtypes. Type III, for example, is divided into four subtypes III-A, B, C and D. While CRISPR-Cas systems of the same subtype encode similar proteins that occupy the same roles, the proteins have often diverged beyond the point of recognition by conventional sequence alignment methods such as Blast, even within a subtype. This level of sequence diversity makes proper identification of the found CRISPR-Cas systems very challenging, and the field has thus far relied upon the gold standard of periodic manual annotations by experts, published once every few years (2, 10, 11). The annotation involves profile HMM searches for finding core genes, followed by the inspection of their neighborhoods, gauging operonic structures, and manual BLAST and PSI-BLAST searches (12). With the increasing number of genome sequences from uncultured microbes and metagenomic data, however, manual annotations cannot keep up and an automated approach is needed, which yields comparable accuracy to manual annotation. Furthermore, research groups working on organisms not yet covered by published annotations have thus far made their own manual annotations, leading to inconsistencies in nomenclature and inaccuracies in some cases.

There have been numerous attempts at devising computational pipelines for the identification of different elements of the CRISPR system, such as CRISPR arrays (13, 14, 15) and CRISPR leaders (16). Detection tools for CRISPR cascades have been implemented only for the simpler Class 2 systems (3, 17, 18, 19), resulting in the detection of novel Class 2 types. These new types have expanded the known diversity and are now being re-engineered for gene-editing applications. Class 1 systems, however, have not been successfully addressed by automated approaches, despite being much more widespread than Class 2 systems. They are difficult to detect due to the signal being spread out across multiple genes, each encoding separate subunits of the effector ribonucleoprotein complex. Previously published methods (20, 21) produce inaccurate results, except for derivatives of the most well-known Class I systems, and not all genes are successfully annotated.

In this work, we present a machine learning (ML) approach intending to capture much of the relevant essence of manual annotation. It is based on evidence for the different Cas proteins to be contained in a specific Cascade, and thus represents genomic CRISPR-Cas systems as cascades of adjacently encoded proteins. These evidences are calculated by newly designed sets of HMM models for each Cas protein, covering the diversity of Cas protein families. The proposed approach solves the problem of classification of new systems into types and subtypes. As our features for the ML approach correspond to evidence for Cas proteins, we can determine Cas proteins whose evidence is critical for predicting a subtype, which corresponds to the concept of signature genes. We show that our approach correctly identifies known signature genes for types and subtypes. In addition, our approach is able provide more information about the composition of cascades. One application is to predict evidence for Cas proteins that have been missed in the Cas protein screening. This provides researchers with hints to search for remote homologs of the missing Cas proteins, or for new proteins that might replace the associated function. Furthermore, we are able to learn association rules, which are subsets of proteins being important to each other, indicating functional modules. As a proof of concept, when we search for Cas proteins associated with an interference protein, our approach finds other interference proteins to be most important. The more interesting cases are undoubtedly with the non-interference protein, where our tool could correctly predict a strong association of the ancillary protein Csn2 with Cas1, consistent with its hypothesized role in adaption. For the helper protein CasR we found that it is associated with different functional modules in subtypes I-A and I-E, indicating a possible functional diversity. Thus, the set of protein associations derived in this manner provides a proper resource for researchers that want to investigate the function of different Cas proteins.

## MATERIALS AND METHODS

### Data collecting and preprocessing

All Cas proteins used in this study were selected from the most recently classified archaeal and bacterial CRISPR-Cas systems (2, 3, 4, 19). We performed an all-against-all sequence similarity comparison on these data using Fasta (22). Subsequently, we clustered the proteins using the Markov Cluster Algorithm (MCL) (23) based on custom similarity criteria (9, 16). These criteria consider the size of the proteins, the length alignment and the relative locations of similar regions between the two compared proteins. After clustering the protein sequences from a specific Cas protein family, we generated a multiple sequence alignment (MSA) using MUSCLE (24). Next, these alignments were converted to HMM profile models by using hmmbuild (25). Except for MCL, all other tools were run with default parameters.

To generate the feature vectors, we ran all HMM profile models using hmmsearch against all cascades. We selected the cascades which had a hit for all proteins annotated for that subtype, and used for the classification pipeline. Cascades that had a missing protein were used instead as a test case for our regression models and the full pipeline.

### Classification of CRISPR Cascades

For this task, we apply ML algorithms onto a finite sample of CRISPR data in order to obtain predictive models that are able to classify protein cascades into their respective subtypes using a data matrix representation (see Results and discussion). Thus, based on the finite sample of data, we investigate the application of classification algorithms that estimate a function which is able to generalize the association between a cascade and its subtypes. As a consequence, we intend to use this function to classify new cascades that were not seen during the training phase into their respective subtypes with a high level of accuracy.

### Prediction of missing Cas proteins

We also investigate the problem of predicting (possibly) missing Cas proteins by estimating their normalized bitscores. For this problem, we modelled it as follows. Given *m* Cas proteins, we filter, for each subtype, its set of *l*<*m* proteins (i.e., all Cas proteins whose bit-score is larger than zero for at least one cascade of the subtype). Next, we induce *l* regressors, where the *j^th^* regressor, *j* ∈{1,⋯,*l*}, predicts the bit-score of the *j^th^* Cas protein using the remaining *l* – 1 proteins as input.

### Experimental evaluation of ML algorithms

We apply three ML algorithms to the preprocessed dataset to obtain predictive models for regression and classification tasks. These algorithms are:

- *Classification and regression trees (CART)* (26), which induces a predictive model represented by a decision tree. This algorithm can induce decision trees for classification (classification trees) and regression (regression trees) tasks. A decision tree is composed by a set of interpretable rules extracted from the training dataset. These rules explain the decisions made by the model to predict the class or regression value for new, previously unseen, examples.
- *Support Vector Machines (SVM)* (27), which induces a binary classifier represented by a hyperplane that separates examples from two classes with the maximum possible separation margin. By using kernel functions, an SVM can be applied to non-linearly separable problems. For multiclass classification tasks, a multiclass dataset is usually first decomposed into several classification binary datasets. SVMs can then be applied to each binary dataset, and their predictions are combined for a multiclasss classification.
- *Extremely Randomized Trees (ERT)* (28), uses an ensemble of decision trees, where each tree is induced using a random subset of the original features. Instead of selecting the best discriminating threshold for each feature considered for a split, as would be the case for classical decision trees, ERT chooses a random threshold value. The final predictions are the average of the predictions of all the decision trees in the ensemble. We can extract the importance of each feature in the classification or regression task from the decision trees in the ensemble. The importance is represented by the decrease in impurity caused by a node that splits the feature, weighted by the number of examples contained in such a node (29), and averaged over all trees of the ensemble.

The model selection and evaluation of predictive models is a widely studied problem in the ML literature. Several works, such as (30, 31, 32), investigate the advantages and drawbacks of different methodologies. Based on these previous studies, we use the nested cross-validation procedure. Given a set of data, the classical cross-validation approach splits the data into *K* mutually exclusive and similar sized subsets called folds. Next, at each iteration, *K* – 1 folds are used for inducing an ML model and the remaining fold for testing it (33, 34). The nested cross-validation approach separates the model selection and evaluation steps, by using two different cross-validation loops: an outer loop, which splits the data into *K*_1_ folds, and is used for model evaluation; and an inner loop, which splits the training data into *K*_2_ folds, and is used for model selection. In this paper, we set *K*_1_ = *K*_2_ = 10, and repeat the evaluation procedure 50 times, due to the variance of the results when considering different splits (32). It is important to mention that, during our experiments, to guarantee that examples from all classes are present in each outer fold, we used only classes containing at least 10 examples.

For each cross-validation iteration, we aggregate the predictions from all folds and calculate a single predictive performance evaluation, in order to avoid any averaging problems that might arise, especially when the dataset is imbalanced (35). For the classification experiments, we used the following evaluation measures: adjusted balanced accuracy score (36, 37), an adaptation of the original accuracy measure that gives higher weights to examples from smaller classes; and the F-score with macro-averaging (38), which is the average F-score among all classes. Both measures treat different subtypes equally. Thus, they do not favor those with the largest numbers of cascades. For the regression experiments, we used the mean absolute error (39), which is the average absolute difference between the expected and the predicted target values.

Regarding the model selection step of each ML algorithm used, we performed a grid search over 20 different hyperparameter combinations, based on the guidelines from the scikit-learn package (40). We describe these hyperparameter grids next. For the CART algorithm, we varied the hyperparameters that determine the maximum depth of the decision tree and the minimum number of examples necessary for a node to become a leaf. For the former, we considered the values in {5,10,15,max}, where max allows the tree to grow as deep as possible. For the latter, we varied the values in {5,6,7,8,9}. For the SVM algorithm, we used a Gaussian kernel, due to its ability to model nonlinear decision boundaries and its reduced number of hyperparameters when compared with another commonly used nonlinear kernel, the polynomial kernel (41). For the cost hyperparameter *C*, we considered the values in {1,10,100,1000}. Regarding the kernel coefficient *γ*, we assessed the values in {0.01,0.1,1,10,100}. Finally, for the ERT algorithm, we varied the ensemble size using the values in {25,50,75,100}, and the quantity of features to be considered when performing a split from the set of values in 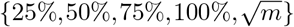, where *m* is the number of known Cas protein families.

### Software and data

CRISPRCasIdentifier is implemented in Python and freely available in http://padilha.github.io/CrisprCasidentifier (password: i4Jpw2hDjPZQi7dBZEIn) under the GNU General Public License v3.0 (GPLv3). It integrates our best ML models for classification and regression in a simple and ready-to-run script. Given an input cascade as a Fasta file, CRISPRCasIdentifier runs our HMM models for the available proteins and labels them according to their best hits. Afterwards, it predicts the normalized bit-scores for the proteins that were not recognized by any HMM. Finally, it proceeds to the classification step, where it is also able to return probabilities for an input cascade belonging to different subtypes.

## RESULTS AND DISCUSSION

### A combined approach to determine Cas proteins and Cascade subtypes

The classification of a subtype is based on the membership for specific Cas proteins. Thus, any ML-based classification of a cascade requires the detection of the contained Cas proteins as a first step. While this first step is commonly performed using Hidden Markov models, a difficulty arises from the fact that a single Cas protein family has to be split into different subfamilies due to the high evolutionary diversity of their members. Due to missing values in the dataset for a family, even the problem of splitting into different subfamilies is not an easy one. Even further, we have observed that the splitting of Cas protein families influence the quality of ML-based subtype classification. This would be quite obvious if subfamilies of individual Cas protein would correlate well with subtypes. The real situation, however, is more complex, partially due to the fact that cascades are composed in a modular way, often involving horizontal gene transfer (2, 9).

In brief and as described in more detail later, our classification approach takes the bit scores for the contained Cas proteins as evidence of their membership to the cascade. We use this information to apply a set of ML algorithms to classify the subtypes of cascades. By generating different divisions of subfamilies for each Cas protein, we obtain different evidences for the contained Cas proteins. Thus, we can investigate which division is best related to subtype evolution. With this holistic view of Cas protein and subtype annotation, we can further examine relations between subtypes and Cas protein membership and as a result re-assure key components of subtytes like signature genes.

### Detection of Cas proteins by families of HMMer models

Our definition of Cas protein subfamilies is based on clustering the known sequences of a specific Cas protein family. We use a set of 3125 cascades with 68594 labeled Cas proteins as a database, and applied different cluster criteria. Each cluster characterizes a subfamily, which is afterwards represented by a HMM model. All models for a Cas protein are grouped, and the best matching HMM for each Cas protein is used to score a new sequence. To cluster the sequences, we performed an all-against-all sequence similarity comparison. Subsequently, we applied the Markov Cluster Algorithm (MCL) (23) to cluster the known sequences for a specific Cas protein family according to their sequence similarities. However, protein sequences can be clustered in different ways, depending on the cut-off for sequence similarity and the requested coverage of the alignment between two sequences. In addition, different hyperparameters for the MCL clustering algorithm result in different data partitions. Each partition defines different subfamilies, for which we train HMM models.

The different clustering approaches thus result in HMM models for different subfamilies, with varying specificity and sensitivity to detect members of a Cas protein family. We created five different collections of HMM models labeled HMM_1_ … HMM_5_ using different hyperparameter values for the clustering algorithm and distinct threshold values for the all-against-all sequence similarity detection (see Methods for detail; the number of models for each Cas protein family is listed in Supplementary Table S1). For a given Cas protein sequence, we applied all HMM models that are contained in a specific collection for that protein family and took the maximum bit score, and zero otherwise. Non-zero values indicate that the investigated protein sequence belongs to the Cas protein family defined by the HMMer model set.

We used different measurements to assess the quality of a specific division represented by a set HMM_*i*_. One quality criteria for a set HMM_i_ is clearly the capability for detecting known members of Cas proteins. Table 1 shows the sensitivity for the five sets HMM_1_ … HMM_5_ by reporting the number of cascades found in each subtype. It is easy to see that the more fine grained sets, HMM_1_, HMM_2_ and HMM_3_, clearly detect more Cas proteins than the less fine grained sets HMM_4_ and HMM_5_.

**Table 1.** Properties and Quality Measurements for the collections HMM_1_ … HMM_5_. (a) Sensitivity of set HMM_*i*_ in detecting Cas proteins, measured by the number of cascades found per subtype. Sets HMM_1_, HMM_2_ and HMM_3_ are more fine grained than sets HMM_4_ and HMM_5_, which detect less Cas proteins overall. (b) Median Accuracy for the classification of subtypes when using set HMM_*i*_ with different ML-approaches to determine the evidence for a Cas protein in a cascade. The quality difference is much lower in the overall task of subtype classification compared to the task of detecting individual Cas proteins.

|  |  | HMM<sub>1</sub> | HMM<sub>2</sub> | HMM<sub>3</sub> | HMM<sub>4</sub> | HMM<sub>5</sub> |
| --- | --- | --- | --- | --- | --- | --- |
| No. models |  | 379 | 385 | 416 | 209 | 201 |
| No. sequences |  | 14674 | 14674 | 23622 | 16018 | 16018 |
| (a) Sensitivity per Subtype | I-A | 116 | 116 | 117 | 0 | 0 |
|  | I-B | 714 | 714 | 707 | 415 | 415 |
|  | I-C | 627 | 627 | 627 | 608 | 608 |
|  | I-D | 138 | 138 | 137 | 100 | 100 |
|  | I-E | 1100 | 1100 | 1102 | 1052 | 1052 |
|  | I-F | 354 | 354 | 353 | 339 | 339 |
|  | I-U | 136 | 136 | 82 | 8 | 8 |
|  | II-A | 319 | 319 | 329 | 244 | 244 |
|  | II-B | 24 | 24 | 35 | 35 | 35 |
|  | II-C | 327 | 327 | 333 | 328 | 328 |
|  | III-A | 374 | 374 | 358 | 321 | 321 |
|  | III-B | 290 | 290 | 288 | 172 | 172 |
|  | III-C | 93 | 93 | 93 | 83 | 83 |
|  | III-D | 181 | 181 | 183 | 45 | 45 |
|  | IV-A | 36 | 36 | 36 | 43 | 43 |
|  | V-A | 18 | 18 | 32 | 27 | 27 |
|  | VI-A | 6 | 6 | 4 | 6 | 6 |
|  | VI-B | 40 | 40 | 40 | 40 | 40 |
| Total Sens. |  | 4893 | 4893 | 4856 | 3868 | 3868 |
| (b) Accuracy | ERT | 0.9908 | 0.9900 | 0.9849 | **0.9916** | 0.9915 |
|  | CART | 0.9690 | **0.9695** | 0.9644 | 0.9541 | 0.9544 |
|  | SVM | **0.9850** | 0.9857 | 0.9802 | 0.9846 | 0.9846 |

In our holistic view of Cas protein detection and subtype classification, however, we want to understand also how the division into subfamilies relates to the cascade subtype, and thus influence the subtype classification. For that reason, we show in Table 1 also as another quality criteria the median accuracy for correctly predicting the subtype of a cascade when using the HMM_*i*_ in a ML-based subtype classification approach as described in the next section. The surprising result is that the sensitivity of a specific set HMM_*i*_ in detecting Cas proteins does not correlate with the accuracy that is achieved in a subtype classification using this set HMM_*i*_.

### A pipeline for CRISPR cascades classification based on Cas protein evidences

Our classification pipeline for CRISPR cascade is described in Figure 1 and has five steps. For each set HMM_1_ … HMM_5_, we build a data matrix for classification and regression analysis of cascades as follows. Usually, a CRISPR cascade 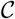 is a collection of Cas proteins and is thus defined as a subset of all known Cas proteins 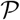 (i.e, 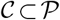. However, when predicting Cas proteins with HMMer models, this would imply a discretization of the bit score that would omit the information about the *evidence* we have for the prediction. For this reason, we define for each cascade 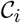 a real vector X_*i*_ of length *m*, where *m* is the number of known Cas protein families, containing an entry for each possible Cas protein.

**Figure 1.**
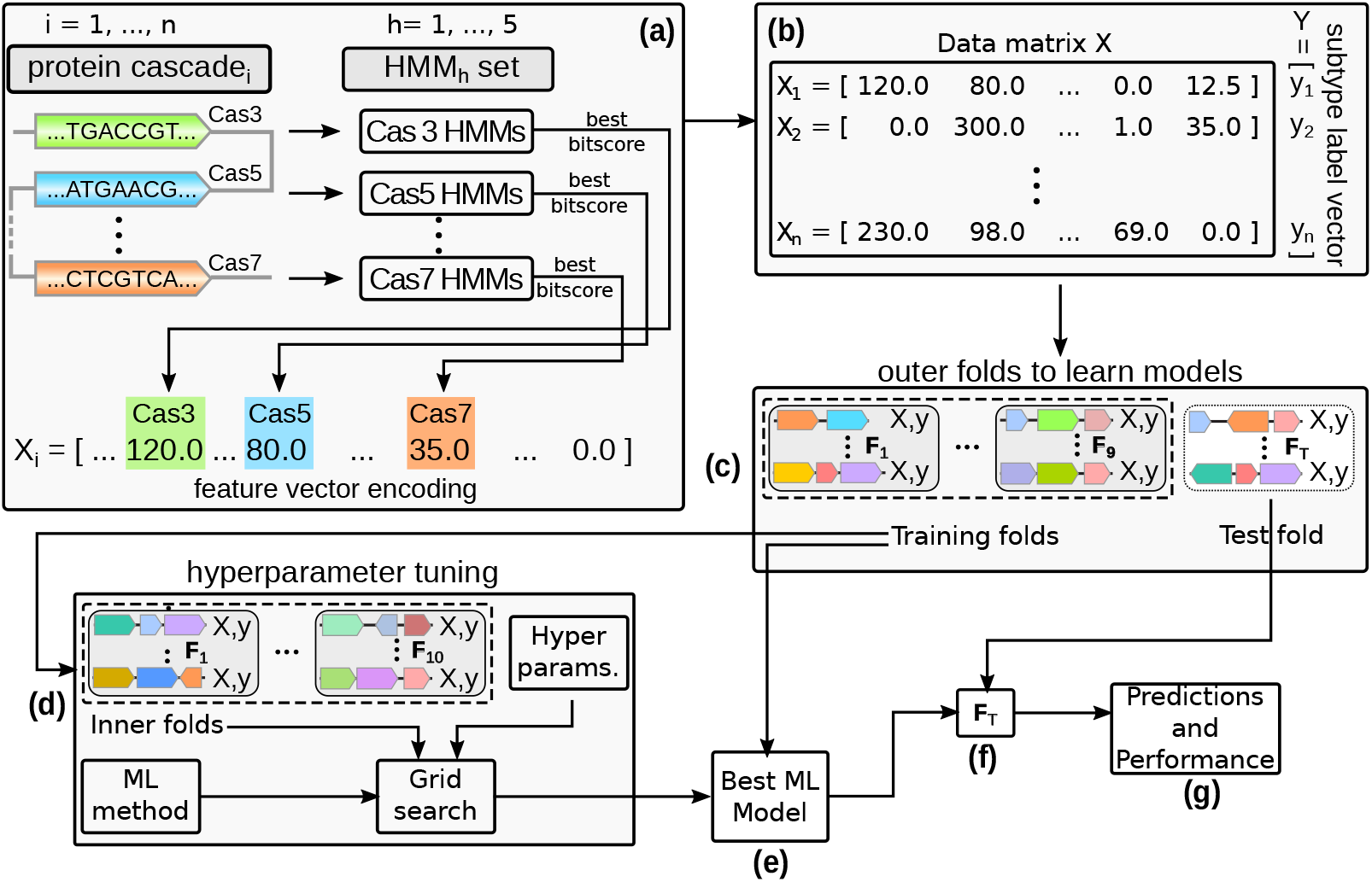
Experimental methodology adopted for this study. (a) Every cascade from our positive set is encoded into a feature vector, which has an entry for each Cas protein family. Given a specific cascade with known Cas proteins, we apply to each Cas protein sequence all HMMs from the set of HMMs that were generated for that specific Cas protein. The best bit-score is included into the feature vector X_i_ encoding the *i^th^* cascade. (b) This feature vector is stored in the data matrix X, together with the known subtype. (c) As the trained model highly depends on the collection of used cascades, we use the ten-fold cross-validation strategy. Thus, we split the training set into 10 subsets called folds. We perform 10 runs, where, in each run, one of the folds is used for testing and the remaining 9 for selecting and training the best ML model. (d) For selecting the best ML model, a similar cross-validation strategy is applied to tune twenty hyperparameter combinations that affect the model predictive performance. Then, in (e), the selected model is trained using the whole training set. Finally, in (f) and (g) we apply the trained model to the respective test set of the outer fold and evaluate its performance.

Each element *X*_ij_ is defined as the best bit-score obtained by 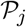 among all HMM models of its family if it is detectable by the models, and zero otherwise (Figure 1a). By concatenating the vectors obtained for all the *n* available cascades, we obtain adatamatrix 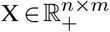 (Figure 1b). In addition, each cascade is associated to a label that indicates its subtype, according to the classification provided by (2, 4, 9).

This data matrix, along with the feature vectors and the subtype labels for all known cascades, is our training data for the subtype classification task. For the evaluation of our classification models, we apply a ten-fold cross-validation procedure on this data matrix. For this, we randomly split the data matrix X into 10 folds (Figure 1c), each one containing a subset of cascades encoded by the associated feature vector. Each vector is annotated (labeled) by its true subtype. For model selection, we perform hyperparameter tuning by employing a grid search over 20 hyperparameter combinations and applying an inner cross-validation loop (Figure 1d, see Methods for detail). After selecting and training the best model (Figure 1e), we have a classifier that, along with a feature vector with HMM bit scores for all known Cas protein families, predicts the subtype of new cascades (Figure 1f and Figure 1g).

### The classification pipeline successfully predicts the subtype of cascades

To evaluate the pipeline, we first assessed whether it can successfully perform the classification task, i.e., correctly predict the subtype of a cascade. As shown in Figure 2 for HMM_1_, the predictive performances, measured by the adjusted balanced accuracy, for CART, ERT and SVM algorithms are above 96%. These high values suggest that, though imbalanced, the cascade subtypes are well-defined in the feature space. It is important to mention that not all cascades are complete in the investigated datasets. Some cascades are composed only by subsets of the Cas proteins that integrate its subtype definition. In Supplementary Table S2, we summarize the percentage of cascades that are complete for each subtype, ignoring Cas proteins that are contained in less than 5% of the cascades of each subtype. We observed in the experimental results that, even though some incomplete cascades are present, the three classifiers were still able to capture the relations among the remaining proteins. The results for the other four sets of HMM models, and for the F-score with macro averaging measure, were similar and allowed us to draw similar conclusions (see Supplementary Figure S1).

**Figure 2.**
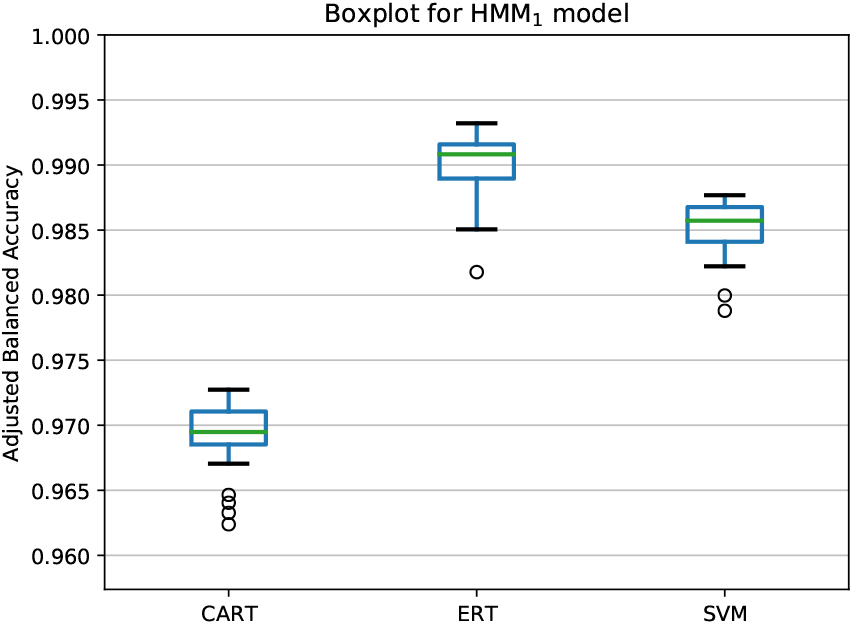
Adjusted balanced accuracy obtained for the 50 repetitions of nested ten-fold cross-validation applying ML algorithms to the dataset generated by the HMM_1_ set. The *x*-axis corresponds to the classifiers induced by different ML algorithms. The *y*-axis shows the range of adjusted balanced accuracy values.

To investigate the prediction quality for specific subtypes, we performed an experiment using the *one-vs-the-rest* strategy (33). Given *k* different classes, the *one-vs-the-rest* strategy induces k classifiers, one for each subtype, which learns how to discriminate this subtype (positive class) from the remaining classes (negative class). In Table 2 we show the average F-scores, after 50 cross-validation repetitions, obtained by the classifiers using the *one-vs-the-rest* strategy. It is clearly visible that the *k* classifiers were able to discriminate each class with a high predictive performance, in agreement with our previous results. In the case of SVM, one can use the margin separating positive and negative data as an additional quality criterion. Again one can see here a clear separation of SVM scores for the positive and negative classes (see Supplementary Figure S2).

**Table 2.**
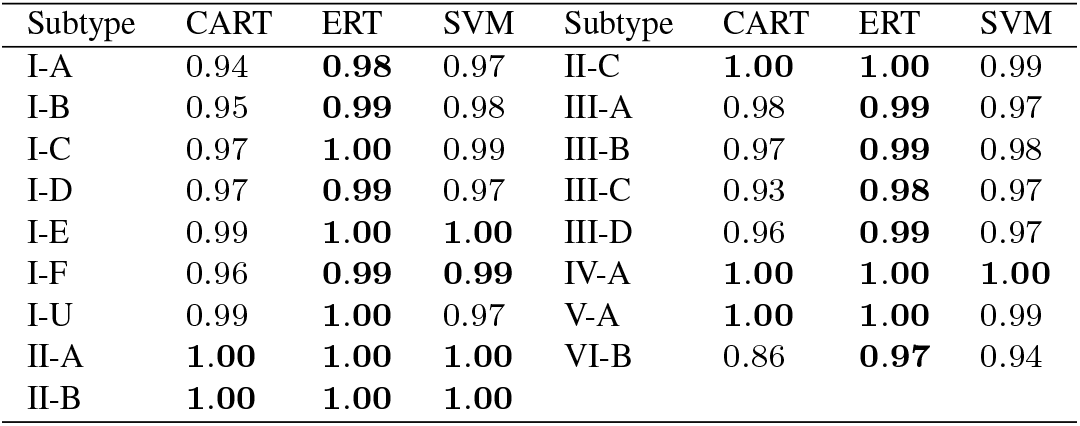
Mean F-scores for 50 nested cross-validation repetitions using the *one-vs-the-rest* strategy and Cas protein set HMM_1_.

### The classification pipeline detects signature proteins

Makarova et *al*. (2) define the presence of unique signature Cas proteins that characterize most of the investigated CRISPR subtypes. According to the authors, signatures usually consist of either one or multiple Cas proteins that co-occur in the same cascade. Based on the aforementioned results, we hypothesize that the classifiers were able to learn these signature proteins. Since *one-vs-the-rest* classifiers introduced in the last section learned how to discriminate a different subtype, we assessed whether it is possible to derive insights about signature proteins for each class by analyzing each classifier separately.

We thus propose a new approach to detect signature proteins for a subtype by determining the importance of a specific feature (i.e, the evidence for a Cas protein in a cascade) to correctly predict the subtype in the respective *one-vs-the-rest* classifier. The rational is that Cas proteins which are highly important for discriminating a specific subtype against all others are likely signature proteins for this subtype. Figure 3 shows the importance of each Cas protein (see Methods for definition of feature importance) in predicting subtype I-D. As can be seen, the importance is specifically high for Cas10(d) (resp. Cas3), which is the signature protein for the subtype I-D (resp. the type I) according to (2). Overall, we observed that Cas10 and Cas3 account, on average, for more than 60% of the importance for classifying I-D.

**Figure 3.**
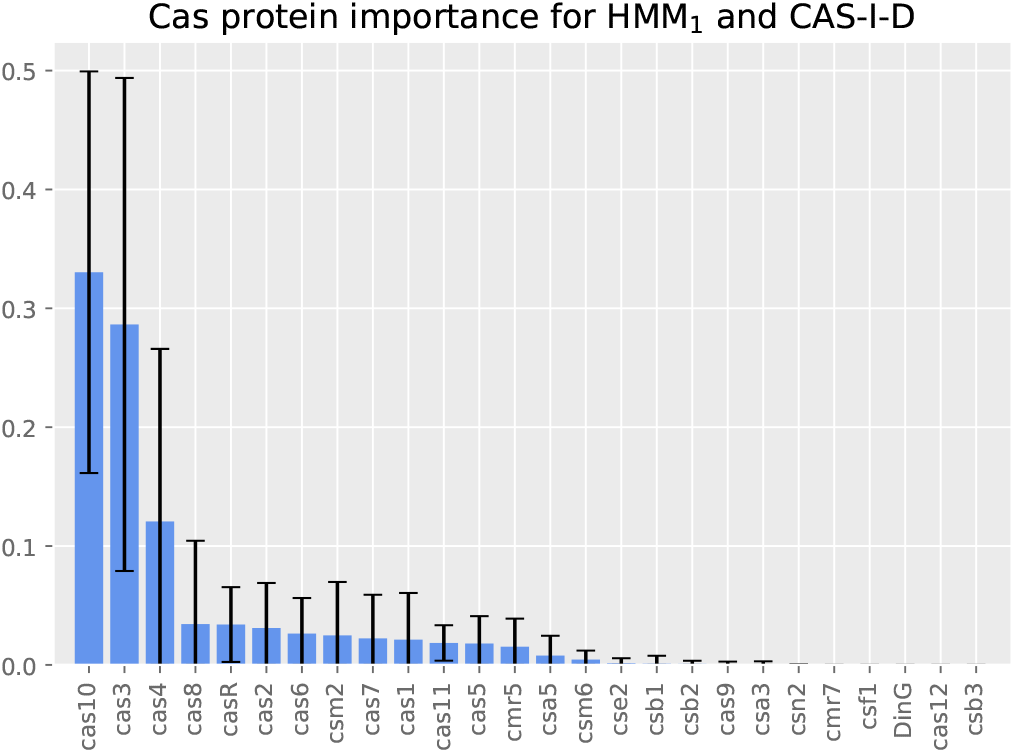
*One-vs-the-rest* average Cas protein importance of ERT for I-D subtype. The *x*-axis presents different Cas proteins. The *y*-axis shows the importance of each Cas protein regarding the decision trees of the ensemble split. Note that the feature importance is not only related with the classification into the I-D subtype, but may also be related to its contribution to classify a cascade into any other subtype. Thus, some of the proteins in the figure may not be related to I-D, but to any other subtype.

To investigate the relation between the two signature genes for proteins Cas10 and Cas3 in more detail, we selected the decision tree obtained by CART for the I-D subtype (Figure 4). In this tree, terminal nodes with the blue color indicate ID classification (positive class), while those with brown color indicate any other subtype classification (negative class). As shown in Figure 4, Cas10 is the most important protein for identifying I-D, which is in agreement with (2), where the subtype I-D is characterized by the presence of a variant of the Cas10 protein (instead of a protein from the Cas8 family, which is common for the other I subtypes) and two variants of the Cas3 protein. Interestingly, we need middle to strong evidence for Cas10 and only weak evidence for Cas3. In the case of weak evidence for Cas10, we also need weak evidence for both Cas3 and Cas1 in order to correct the missing 36 examples, albeit in this case the classification would not be pure anymore. Overall, it can be observed that CART was able to correctly model this signature, since most of the nonterminal nodes refer to these proteins, indicating that they are the most important features in this subtype.

**Figure 4.**
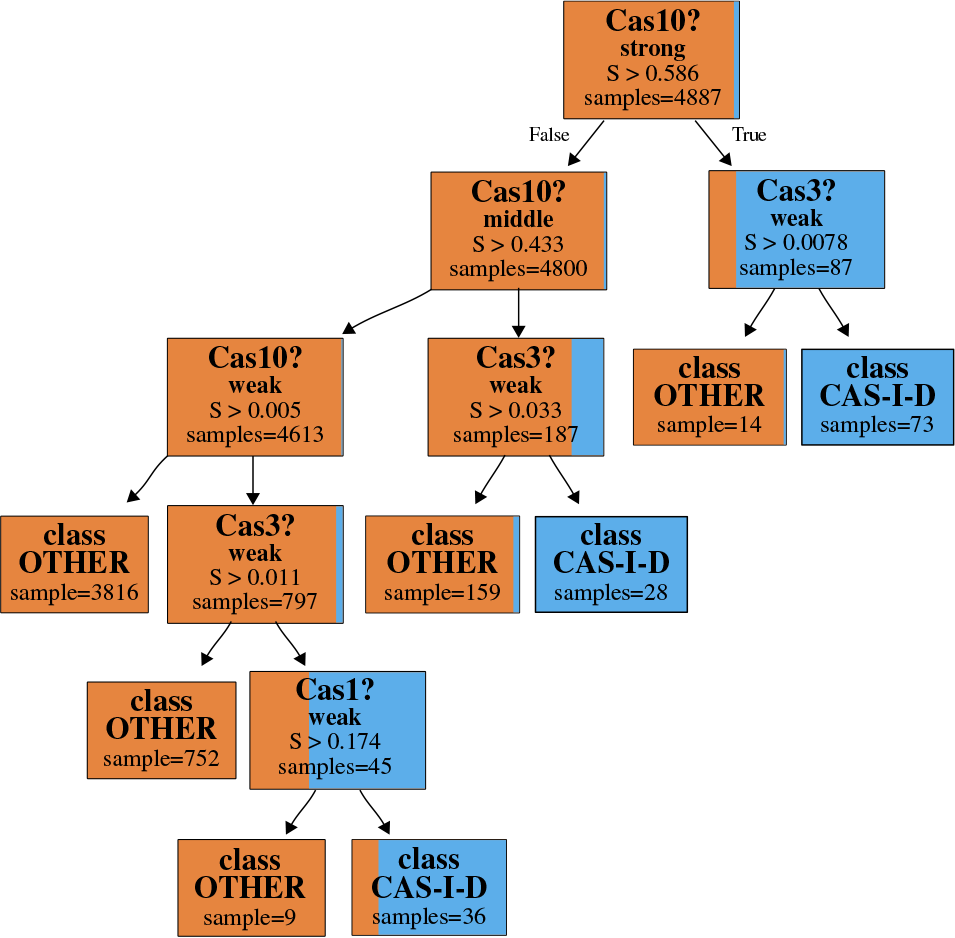
Reduced *one-vs-the-rest* CART for the I-D class (see Supplementary Figure S3 for full tree). Cascades that are labeled as subtype I-D are highlighted in blue, the others in brown. Each node shows the fractions of class I-D and other cascades, indicating the purity of the node. The number of cascades is shown under the “samples” entry. In each node, we query for evidence of a specific Cas protein, indicated by the score calculated by the HMM family models. As one can see, a strong evidence for Cas10 immediately points to a subtype I-D (top node and right branch). Otherwise, if we have middle evidence for Cas10, we need at least weak evidence for Cas3 to determine subtype I-D. Finally, if we have only weak evidence for Cas10, we need at least weak evidence for Cas3 and also for Cas1 to determine subtype I-D (left branch). However, the classification is not pure anymore (bottom nodes).

Since the current classification (2) is based only on the interference module, the adaptation-related Cas proteins (Cas1, Cas2 and Cas4) should not have a high importance for our classification pipeline. Thus, in another experiment, we removed these proteins and the process proteins (Cas6), and tested the predictive performance of our classification pipeline when removing this information. The obtained results were similar to those previously discussed in this section and support our discussion and main conclusions (see Supplementary Figure S4), strengthening the hypothesis that our ML-based approach captured biologically relevant information.

All the aforementioned examples illustrate how our ML models are able to learn the protein signatures without any extra information other than the normalized bit scores and cascade subtype labels. These results validate our hypothesis and provide models that are able to automatically categorize new cascades with a high predictive accuracy.

### Regression instead of classification learns association rules

In our next set of experiments, we were interested in answering the question of whether some Cas proteins tend to be co-occurring frequently with other proteins. To answer this question, we hypothesized that they form a functional module. However, as we have varying information about the evidence for a specific Cas protein, and there is also some redundancy and flexibility in forming this module, we followed an approach different from that described in the previous section. We believe that if a specific Cas protein is frequently associated with other Cas proteins, it is possible to predict the evidence for this protein by relying only on the known evidence for the other members of the functional module. We can confirm this belief by removing a specific Cas protein from the feature vector, and predicting the “expected” normalized bit score for this protein from the remaining feature vector. This amounts to learning a regression model from known examples.

Association rules can now be inspected by determining again the important features (i.e., Cas proteins) to predict the correct evidence for a specific Cas protein. In Table 3, we list the three most important proteins for some target Cas proteins in some subtypes. In this case, for predicting evidence for Cas10d in subtype I-D, we need the information about Cas3, Cas5 and Cas7. In agreement with the fact that subtypes are mainly associated with the interference complex (2), we find that for the interference proteins Cas10d, Cas3 and Cse2, the associated proteins are also interference proteins. For the noninterference proteins Csn2 and Cas4 in II-A and II-B, not only is Cas9 an interference and signature protein for type II, but it is associated with them as well as the adaptation proteins Cas1 and Cas2. Interestingly, though Cas9 information is important for Cas4, Cas1 is actually more significant for Csn2. This is in agreement with the hypothesized role of Csn2 in the adaptation process (42, 43, 44, 45).

**Table 3.** Top 3 most important proteins according to ERT when trying to predict a target protein across different subtypes. For the interference proteins Cas10d, Cas3 and Cse2, the other most important Cas proteins are also interference proteins. For non-interference proteins, other Cas proteins linked to adaptation, e.g. Cas1 and Cas2, are also important. The helper protein CasR seems to have different modules associated in I-A and I-E.

| Subtype | Target protein | Most important proteins |
| --- | --- | --- |
| I-D | Cas10d | (Cas3, 0.37), (Cas5, 0.16), (Cas7, 0.12) |
| I-D | Cas3 | (Cas11, 0.41), (Cas10, 0.28), (Cas5, 0.14) |
| I-E | Cse2 | (Cas8, 0.23), (Cas5, 0.22), (Cas7, 0.18) |
| II-A | Csn2 | (Cas1, 0.57), (Cas9, 0.24), (Cas2, 0.19) |
| II-B | Cas4 | (Cas9, 0.84), (Cas2, 0.12), (Cas1, 0.04) |
| I-A | CasR | (Csa5, 0.33), (Cas5, 0.17), (Cas6, 0.15) |
| I-E | CasR | (Cas1, 0.38), (Cas8, 0.27), (Cas7, 0.17) |

An interesting case to consider is CasR, a helper protein of unclear function. This protein seems to have different roles in subtypes I-A and I-E, and also appears to be associated with the different proteins in I-A and I-E (see Table 3, last two rows). In I-A, the most important proteins are Csa5, Cas5 and Cas6, whereas in I-E they are Cas1, Cas8 and Cas7.

### The ML-approach can handle missing Cas proteins

In real world applications, it is necessary to predict the subtype for cascades that have one or more Cas proteins missing, i.e., there are no hits in their corresponding HMM models during the preprocessing step (Figure 1a). Therefore, it is important to assess whether our ML-based pipeline can handle these cases, which are frequently occurring in real application scenarios. We furthermore assessed whether we can also predict the missing evidence for these proteins.

Concerning the first task, we took cascades that had a missing protein in the annotation. Note that we had excluded these cascades before training the regressor (see Methods), thus forming an independent test set. We then applied our classifiers to predict the subtype for these incomplete cascades. Table 4 shows the classification results for all ML algorithms on this independent test. Especially the ERT-based classifier is still capable to predict the corrected subtype with high quality, even in the hard case of incomplete annotation.

**Table 4.** Average adjusted balanced accuracy for classification on the independent test set, consisting of cascades with one Cas protein missing.

| Classif. | HMM<sub>1</sub> | HMM<sub>2</sub> | HMM<sub>3</sub> | HMM<sub>4</sub> | HMM<sub>5</sub> |
| --- | --- | --- | --- | --- | --- |
| CART | 0.53 | 0.53 | **0.69** | 0.50 | 0.50 |
| ERT | 0.72 | 0.72 | **0.76** | 0.72 | 0.72 |
| SVM | 0.58 | 0.58 | **0.72** | 0.62 | 0.62 |

One can also see that in this real world scenario, the more fine grained Cas protein set HMM_3_ is clearly outperforming the other HMM collections.

In our experiments, we observed that most of the aforementioned cascades contained only one protein that did not achieve any hit for the HMM models of its family. We thus investigated whether we could predict the missing evidence using the previously described regression approach, trained on all subtypes. The basic idea is that a high predicted evidence for a missing protein is a hint for researchers to perform an in-depth attempt to annotate the missing protein, or to search for new proteins that might replace the function of the missing protein.

To investigate this, we removed one bit score for a specific protein, and learn a model that is able to predict this bit score using the evidence information from the remaining proteins. In Figure 5, we present the Cas protein regression results for ERT; the regressor with the lowest mean absolute error for subtypes I-A and I-E in the dataset generated by HMM_1_. Concerning the different subtypes and datasets, the other results were similar and are presented in our Supplementary material, Figures S5–S10. We can see that the missing proteins are well predicted in general. The core proteins Cas1 … Cas8 are especially well predicted by the approach, showing a high interdependence between these core proteins and other Cas proteins important for the subtype. We also observed that for proteins different from the core Cas proteins such as CasR, the size of the data basis (i.e., number of known cascades) for the subtype is influencing the prediction quality, indicating a more variable (or complex) interaction between these proteins and other proteins important for the subtype.

**Figure 5.**
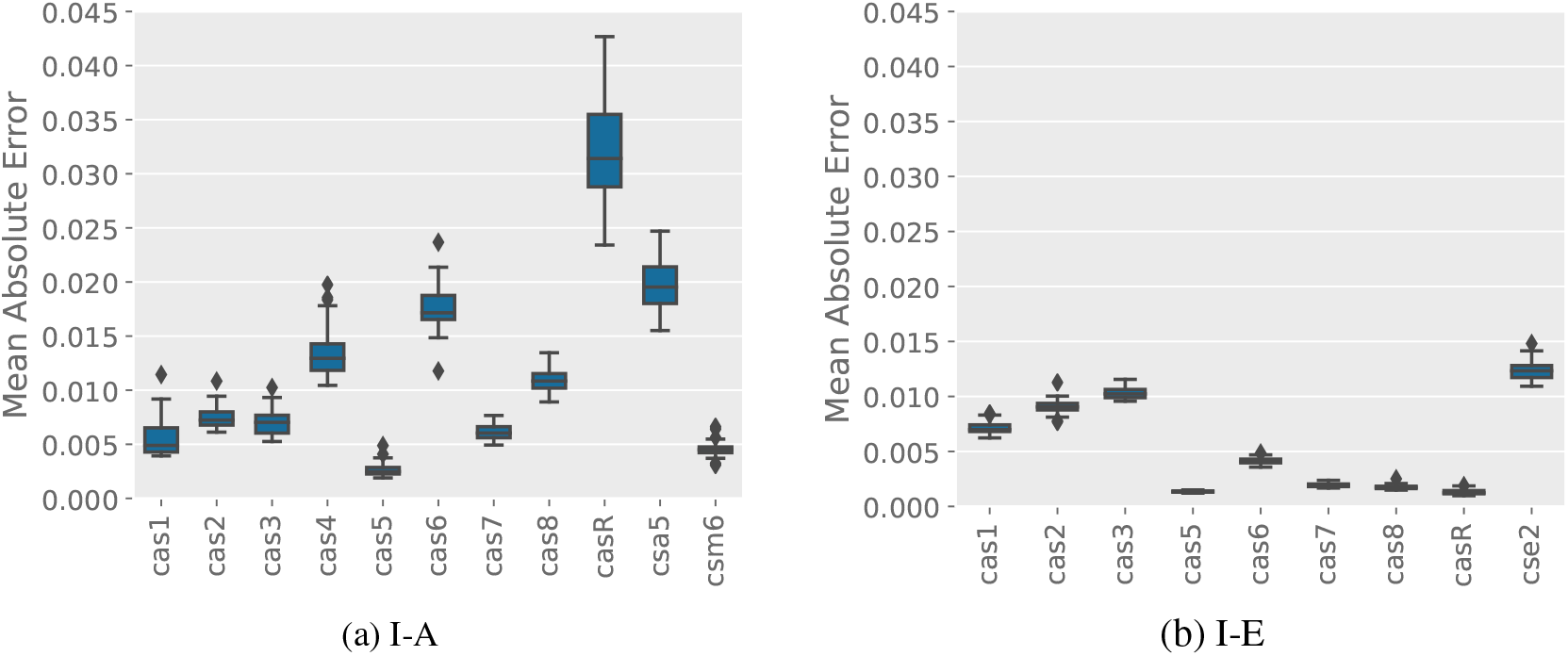
Mean absolute error results for Cas proteins contained in I-A (a) and I-E (b) subtypes over 50 nested cross-validation repetitions. In the *x*-axis we list the different Cas proteins that were used as target variables. In the *y*-axis we present the mean absolute error values. In general, missing proteins are well predicted, especially in the case of the core Cas proteins Cas1 to Cas7. For other Cas proteins like CasR, the prediction quality is varying between I-A and I-E. This is likely to the higher amount of I-E cascades in the data basis, indicating a more complex relationship between CasR and other Cas proteins.

We also observed that the ERT obtained in general the best results for Cas protein regression (see Supplementary Figures S5–S10). In most cases, ERT achieved mean absolute error values below 0.05 for the normalized bit score prediction. These results confirm the relevance of building specific regressors for each Cas protein inside of a specific subtype for the identification of unknown or possibly missing Cas proteins, when the label of the cascade of interest is known.

Finally, as a proof of concept, we looked at the independent test case consisting of cascades with one missing protein, and applied our regression approach in order to determine the cascades where the evidence for having a specific missing protein was predicted to be high. These cases would be good candidates for missing annotations. We found 13 cascades that predicted a missing DinG protein of which three had a predicted evidence of 0.5 or higher. By applying a HHblits (46) search for all ORFs in the respective genome of these three cascades, we found an ORF with convincing homology to DinG-proteins in each case (see Figure 6A for an example). Another case was Cas2, where we found 13 cascades with a missing Cas2 protein predicted. We again used HHblits on all ORFs in the genome of the top three cascades, and found one case with a convincing Cas2 homology (see Figure 6B).

**Figure 6.**
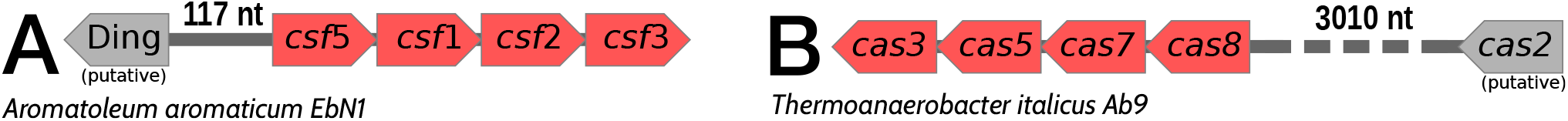
The Cascades with missing proteins. A) In this genome, we predicted a DinG protein missing in the cascade with evidence > 0.5. The HHblits (46) search in this genome for all ORFs determined one ORF 117nt upstream of the cascade with a high confidence score for a DinG homology (E-value: 7.6e-22). B) In the case of Cas2, the predicted evidence was lower, between 0.221 to 0.165. Nevertheless, we found one ORF with a high confidence score for Cas2 homology (E-value: 1e-37) 3010nt downstream of the cascade.

## CONCLUSIONS

In this paper we introduced a new ML-based pipeline for the identification and classification of genomic CRISPR-Cas systems. To assess the predictive performance of this approach, we conducted an in-depth investigation of the suitability of commonly used ML algorithms for the task by using the normalized profile HMM search bit scores of Cas proteins as input, and classifying cascades of proteins to their respective subtypes according to the most recent classification (2, 4, 9).

Overall, this work covers three different research issues: (i) the classification of cascades; (ii) the prediction of normalized bit scores for missing Cas proteins; and (iii) the investigation of the properties of CRISPR types and subtypes. Concerning topic (i), our classification models were able to achieve very high classification performance, above 0.96, in terms of the adjusted balanced accuracy score. Thus, they are well placed for the prediction of CRISPR systems of newly sequenced organisms, or metagenomic data with sufficient read length to cover the full cascade in one contig. In addition, we introduced a new method for determining signature genes, which are genes most important for predicting the correct subtype. This approach was able to properly learn the known signature genes of CRISPR-Cas subtypes without any extra information other than the available protein cascades and their labels, but provides additional information about the composition of cascades. In topic (ii), our regressor models achieved very small deviations between the expected and predicted normalized bit scores for different Cas proteins across the different subtypes. This illustrates the usefulness of these regressors on new cascades that have missing hits for some Cas proteins. A high bit score provides a hint to researchers to search for more diverged forms of the protein, or to look for proteins which could replace the missing function. The analysis performed under topic (iii) enabled us to correctly identify known signature genes, and to identify putative functional modules. Overall, it provided us with a set of association rules for potential use in more advanced classification scenarios, in addition to providing insights about the biology of the systems.

Manual annotation is the gold standard when it comes to classification and identification of genomic CRISPR-Cas systems. Replacing or supporting this process with an automated algorithm requires a degree of flexibility that is challenging to model. CRISPRCasIdentifier provides a boost in classification accuracy when compared to existing tools, because it builds on an understanding of the manual annotation process. We made CRISPRCasIdentifier available for researchers to use with their own data.

